# Implantation failure linked to altered uterine NK cells in PCOS-like mice

**DOI:** 10.64898/2026.02.12.705498

**Authors:** Sara Torstensson, Haojiang Lu, Allan Zhao, Camille Gauthier, Gustaw Eriksson, Eva Lindgren, Qiaolin Deng, Maria H. Johansson, Benedict J. Chambers, Anna Benrick, Elisabet Stener-Victorin

**Affiliations:** Department of Physiology and Pharmacology, Karolinska Institutet, 171 77 Stockholm, Sweden; Department of Microbiology, Tumor and Cell Biology, Karolinska Institutet, 171 77 Stockholm, Sweden; Department of Medicine Huddinge, Karolinska Institutet, 141 52 Huddinge, Sweden; Department of Physiology, Institute of Neuroscience and Physiology, Sahlgrenska Academy, University of Gothenburg, Box 432, 40530 Gothenburg, Sweden

**Author notes:** **Corresponding author** Elisabet Stener-Victorin, Department of Physiology and Pharmacology, Karolinska Institutet, 171 77 Stockholm, Sweden.

**Keywords:** Polycystic ovary syndrome, uterine NK cells, hyperandrogenism, reproductive immunology

## Abstract

Uterine NK (uNK) cells are essential for reproductive function, yet little is known how they are affected in polycystic ovary syndrome (PCOS), despite the strong association between PCOS and reproductive complications. We demonstrate that implantation failure coincides with distinct phenotypic alterations of uNK cells in a PCOS-like mouse model. Hyperandrogenism caused an increased influx of conventional NK (cNK) cells into the uterus, which contributed to an augmented uNK cell population, while tissue-resident NK (trNK) cells remained unchanged. Notably, CD69^+^ trNK cells were reduced and seemingly compensated by an upregulated expression of CD69 on cNK cells in the uterus. The maturation of uNK cells was disrupted and plausibly linked to an inability of cNK cells to convert into trNK cells. The inhibited maturation was associated with a reduced expression of the inhibitory receptor NKG2A, demonstrating an impaired education of uNK cells. This disruption of uNK cells could contribute to endometrial dysfunction and may be an underlying factor to reproductive comorbidities such as implantation failure, miscarriage and pre-eclampsia in PCOS.

## Introduction

Polycystic ovary syndrome (PCOS) is strongly associated with reduced fertility, early miscarriage, and preeclampsia^1,2^. However, the impact of hyperandrogenism, a key feature of PCOS, on uterine natural killer (uNK) cells remains poorly understood, despite their critical role in reproductive function. Notably, uNK cell dysfunction has been implicated in several pregnancy complications that overlap with common comorbidities in PCOS^3–7^.

uNK cells, defined as NK cells residing in the uterus, are functionally distinct from conventional NK (cNK) cells, exhibiting lower cytotoxic potential^8,9^. Instead, uNK cells are crucial for the formation of spiral arteries^10–15^ and are believed to mediate implantation through interactions with the trophoblast^11,16^. The importance of uNK cells for successful implantation and a healthy pregnancy is underscored by studies demonstrating that incompatibilities between maternal human killer-cell immunoglobulin-like receptors (KIRs) haplotypes and fetal KIR ligands predispose individuals to pre-eclampsia and adverse pregnancy outcomes^3,6,17^.

The majority of uNK cells are tissue-resident, discriminated by their expression of the integrin receptor CD49a in mice^18^. The origin of uNK cells remain elusive, with some reports indicating local proliferation of tissue-resident uNK cells^19^ and others demonstrating an influx of NK cells from the periphery^20^. These theories are not mutually exclusive, and recent work has shown that cNK cells can transform into tissue-resident NK (trNK) cells in response to murine cytomegalovirus infection^21,22^. Furthermore, autocrine signaling of transforming growth factor-β1 (TGF-β1) drives this conversion and is also required for the development and maintenance of trNK cells in the uterus^22^. The transformation of cNK cells into trNK cells is also under the regulation of progesterone in the cycling uterus of mice^23^.

Many reproductive complications associated with PCOS are believed to result from endometrial dysfunction, which is characterized by dysregulated enzymes involved in sex hormone production, as well as altered cell signaling, metabolic pathways, and expression of sex hormone receptors^24^. Additionally, metabolic comorbidities such as obesity and type 2 diabetes are common among women with PCOS^25–27^ and are known to negatively impact reproductive function^28–31^. How the uterine environment in PCOS affects immune function remains largely unknown. Single-cell analysis of endometrium from women with PCOS has revealed changes in both composition and gene expression of immune cell populations during the proliferative phase^32^. We have also previously reported that androgen exposure causes tissue-specific alterations of immune populations in a well-established mouse model of PCOS, with a pronounced increase in NK cells in the uterus^33^.

Disruption of uNK cells may underlie the endometrial dysfunction observed in PCOS. To elucidate this, we investigate how hyperandrogenism alters the phenotype and functional profile of uNK cells, and how these changes relate to reproductive outcomes in a peripubertal dihydrotestosterone (DHT)-exposed mouse model that recapitulates key features of PCOS.

## Results

### Augmented population of uNK cells in PCOS-like mice is due to an increased infiltration of cNK cells

We have previously reported that NK cells are increased in the uterus of peripubertal DHT-exposed PCOS-like mice^33^. To determine whether the augmented population of uNK cells stems from increased local proliferation of tissue-resident NK cells (trNK) or from elevated infiltration of conventional NK cells (cNK), we analyzed phenotypic markers of uNK cells from peripubertal DHT-exposed PCOS-like mice by flow cytometry (Figure 1a). The markers CD49a and CD62L were used to distinguish trNK cells from cNK cells that recently migrated from peripheral blood. An overall increased frequency of NK cells (defined as CD45^+^Lin^−^NK1.1^+^, Figure S1a) was observed in the uterus of PCOS-like mice, confirming our previous findings (Figure S1b). As expected, the majority of uNK cells were tissue-resident, defined by their expression of CD49a (Figure 1b), while cNK cells (defined as CD62L^+^ NK cells) constituted a considerably smaller population in the uterus of control mice (mean 15%, Figure 1c). While no change in the frequency of trNK cells was observed, cNK cells were drastically increased in the uterus of PCOS-like mice, indicating a higher influx of cNK from the periphery. This increase was prevented in mice co-treated with the androgen receptor antagonist flutamide (Figure 1c). The notion of an increased infiltration of cNK cells was further supported by unchanged proportions of uNK cells expressing the tissue residency markers CXCR6 and CD69, the latter also a marker of activation on NK cells, and an increased frequency of DX5^+^ NK cells in the uterus of PCOS-like mice (Figure S1c-e). In the analysis of CD49a^+^ and CD62L^+^ uNK cells, it was noticed that NK cells single positive for CD49a or CD62L constituted only about 60-70% of all uNK cells. A substantial proportion of uNK cells expressed neither CD49a nor CD62L (Figure 1d-e). While the proportion of this double-negative (DN) population was unchanged among groups (Figure 1e), it contributed considerably to the higher number of uNK cells in PCOS-like mice (Figure 1f and Figure S1g). A CD49a^−^CD62L^−^ DN population was found in blood too, albeit notably smaller (Figure 1g). While approximately 26% of uNK cells were DN (Figure 1d), this population constituted only 6% of NK cells in circulation. It is therefore unlikely that the DN population seen in the uterus is a result of blood contamination. In contrast, the CD49a^+^CD62L^+^ double-positive (DP) population constituted less than 5% of uNK cells in controls but was noticeably expanded in PCOS-like mice (Figure 1d). Both the DN and DP populations in mice co-treated with flutamide were similar to controls. Collectively, androgen exposure in PCOS-like mice causes an increased influx of cNK cells into the uterus, as well as an increased number of DN uNK cells.

**Figure 1.**
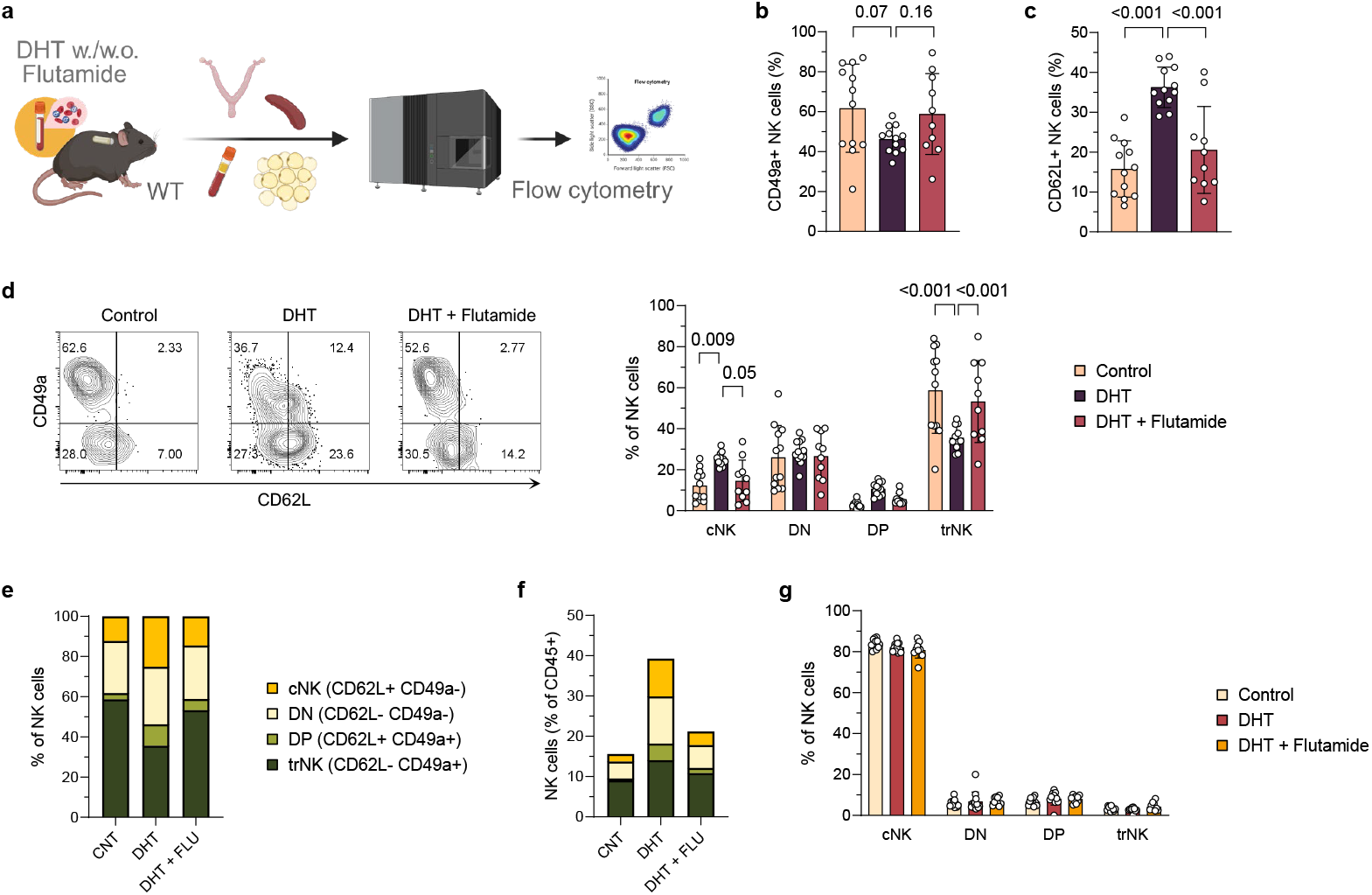
Augmented population of uNK cells in PCOS-like mice is due to an increased infiltration of cNK cells. (**a**) Experimental design. (**b**) Frequency CD49a+ uNK cells (as % of Lin-NK1.1+ cells). (**c**) Frequency CD62L+ uNK cells (as % of Lin-NK1.1+ cells). (**d**) Representative flow cytometry plots and quantification of conventional (cNK; CD62L+CD49a-), double-negative (DN; CD62L-CD49a-), double-positive (DP; CD62L+CD49a+) and tissue-resident (trNK; CD62L-CD49a+) uNK cells. (**e**) Proportions of subsets, out of all uNK cells. (**f**) Proportions of subsets, out of all CD45+ immune cells in uterus. (**g**) Frequency of cNK, DN, DP and trNK cells in blood, expressed as a percent of NK cells. Data are presented as means ± SD.

### Distinct phenotypic alterations of uNK subsets in PCOS-like mice

With a drastic change in both the number and composition of uNK cells in PCOS-like mice, we next examined how the phenotype of these subsets was affected by androgen exposure. First, to better define the CD49a^−^CD62L^−^ DN population of uNK cells, we assessed the expression of tissue residency markers on this subset. While the average proportion of DN uNK cells expressing the tissue residency marker CXCR6 was less than 1% in controls, this was modestly increased in DHT-exposed mice but not in mice co-treated with flutamide (Figure 2a). Overall, CXCR6 expression was primarily restricted to trNK cells (Figure S2a). Similarly, less than 10% of DN uNK cells expressed CD69 in controls (Figure 2b). The low expression of CD69, a marker of both tissue residency and activation on NK cells, and a lack of expression of CXCR6, plausibly indicate a greater similarity of DN uNK cells to cNK cells than to trNK cells in the normal cycling uterus. Surprisingly, the frequency of CD69^+^ DN uNK cells was greatly increased in PCOS-like mice (Figure 2b). Since no difference in the overall frequencies of CD69^+^ uNK cells was observed (Figure S1d), this increase must coincide with a lower expression of CD69 on another subpopulation. Indeed, when specifically analyzing trNK cells, which otherwise are the main CD69-expressing population, a clear decrease in CD69^+^ trNK cells was seen in PCOS-like mice (Figure 2c). Further analysis revealed that a small but distinct proportion of cNK cells in the uterus of PCOS-like mice also expressed CD69, in strong contrast to controls where CD69^+^ cNK cells constituted approximately 1% of the total uNK cells (Figure 2d and Figure S2b). Notably, no change in CD69 expression was observed on NK cells in blood (Figure S2c), indicating that this effect is coupled to the infiltration of cNK cells into the uterine tissue.

**Figure 2.**
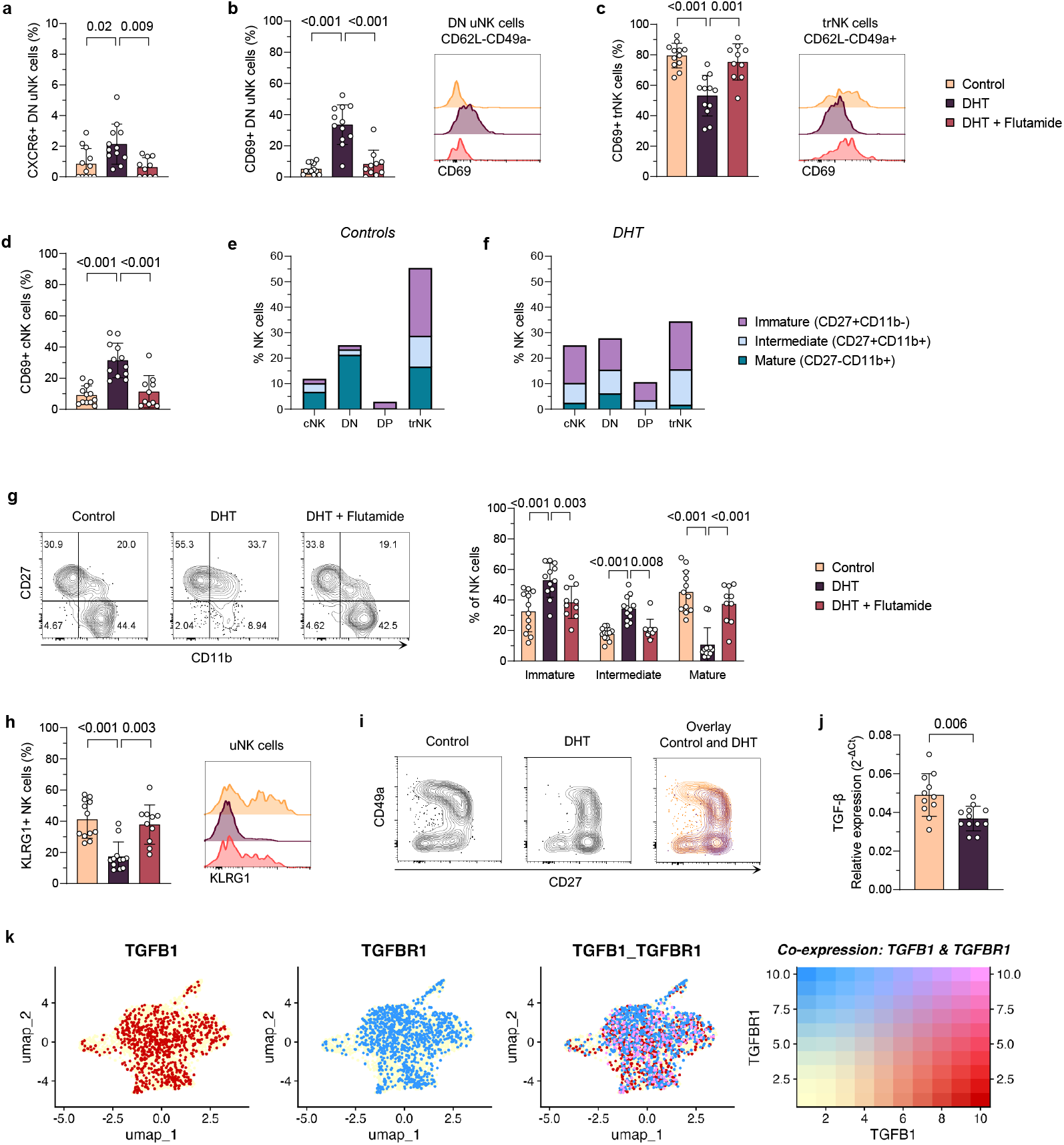
Distinct phenotypic alterations of uNK subsets in PCOS-like mice. (**a**) Frequency CXCR6+ double-negative (DN; CD62L-CD49a-) uNK cells. (**b**) Quantification and representative flow cytometry histograms of CD69+ DN uNK cells. (**c**) Quantification and representative flow cytometry histograms of CD69+ tissue-resident NK (trNK) cells in uterus. (**d**) Frequency CD69+ conventional NK (cNK) cells in uterus. (**e**) Proportions of maturation among subsets in controls, out of NK cells in uterus. (**f**) Proportions of maturation among subsets in DHT-exposed PCOS-like mice, out of NK cells in uterus. (**g**) Representative flow cytometry plots and quantification of immature (CD27+CD11b-), intermediate (CD27+CD11b+) and mature (CD27-CD11b+) uNK cells. (**h**) Quantification and representative flow cytometry histograms of the maturation marker KLRG1+ uNK cells. (**i**) Representative flow cytometry plots of expression pattern of CD49a and CD27 on NK cells from controls compared to DHT-exposed PCOS-like mice. (**j**) Relative expression of TGF-β in uterus. (**k**) UMAP projections of integrated snRNA-seq data of endometrial NK (eNK) cells from women with and without PCOS, displaying expression of TGF-β (TGFB1, red), TGF-β receptor 1 (TGFBR1, blue), and co-expression of TGF-β and TGF-β receptor 1 (purple). Data are presented as means ± SD.

It has been shown that immature cNK cells are more prone to convert into trNK cells when stimulated with TGF-β1 than mature cNK cells^22^. Therefore, the maturation of uNK cells was analyzed among different subpopulations. In the cycling uterus of control mice, a significant proportion of trNK cells was immature (∼40%, CD27^+^CD11b^−^), while ∼30% were mature (CD27^−^CD11b^+^) and ∼20% were in an intermediate state (CD27^+^CD11b^+^) (Figure 2e and Figure S2e). In contrast, DN uNK cells were predominantly mature, while the small DP population was exclusively immature (Figure 2e and Figure S2f-g). Among the cNK cells, roughly 50% were mature, followed by an intermediate population that were slightly larger than the immature population (Figure 2e and Figure S2h). The maturation of uNK cells was strikingly altered in PCOS-like mice (Figure 2f). Here, immature and intermediate uNK cells dominated all subpopulations while mature uNK cells were drastically reduced (Figure 2f and Figure S2e-h). This prominent effect on maturation was evident for the whole population of uNK cells and was prevented in mice co-treated with flutamide (Figure 2g). In line with this finding, a decreased frequency of uNK cells expressing the maturation marker KLRG1 was seen in PCOS-like mice (Figure 2h and Figure S2i). When analyzing flow cytometry data, a plausible association between maturation and tissue residency was noted in the expression pattern of CD27 in relation to CD49a (Figure 2i and Figure S2j). This pattern appears to differ between controls and PCOS-like mice (Figure 2i), which could suggest that immature cNK cells fail to transition into mature trNK cells.

Autocrine TGF-β1 signaling is essential for the conversion of cNK cells into trNK cells^22^. To explore whether the increased influx of immature cNK cells could be linked to impaired differentiation into trNK cells in PCOS-like mice, the expression of TGF-β1 was assessed. A significantly lower expression of TGF-β1, measured by qPCR, was found in the uteri of PCOS-like mice compared to controls (Figure 2j), which could support an impaired differentiation of cNK cells into trNK cells. In humans, decidual stromal cells and NK cells from peripheral blood has been shown to produce TGF-β1^22,34^. However, autocrine signaling of TGF-β1 in uNK cells has so far only been demonstrated in mice^22^. Therefore, we utilized our PCOS-endometrial cell atlas (PECA) of human endometrial single nuclei RNA sequencing dataset^32^ to assess the expression of TGF-β1 in endometrial NK (eNK) cells. TGF-β1 was indeed expressed by eNK cells (Figure 2k), and an equal expression was found among the different subsets eNK1, eNK2, and eNK3 (Figure S2k-m). Importantly, co-expression of TGF-β1 and the TGF-β1 receptor (TGFBR1) was observed on all subsets (eNK1-3), demonstrating that eNK cells fulfill the requirement for autocrine signaling.

In summary, uNK subsets have distinct phenotypes and androgen exposure causes specific alterations to these. In particular, the maturation of uNK cells appears to be inhibited in PCOS-like mice, and lower expression of TGF-β1 might link this to impaired differentiation of immature cNK cells into trNK cells.

### Impaired education of uNK cells in PCOS-like mice

Education of uNK cells is a key process to regulate their function and is important for the fetal-maternal environment in healthy pregnancy^15,35^. Therefore, the effect of androgen exposure on the expression of NKG2A, one of the major receptors mediating NK cell education, was examined. A decreased frequency of NKG2A^+^ uNK cells was observed in PCOS-like mice, compared to controls or mice co-treated with flutamide (Figure 3a). The altered education appeared to be tied to maturation, as a smaller proportion of intermediate CD27^+^CD11b^+^ uNK cells expressed NKG2A (Figure 3b). However, since this intermediate population was expanded in PCOS-like mice (Figure 2h), a smaller proportion would not necessarily account for the overall reduction in NKG2A^+^ uNK cells. Indeed, subsequent analysis revealed that this reduction was due to fewer mature NKG2A^+^ uNK cells (Figure 3c and Figure S3b). Although the proportion of mature uNK cells expressing NKG2A was unchanged in PCOS-like mice (Figure 3b), the drastic decrease in mature uNK cells (Figure 2g) significantly affected the overall frequency of NKG2A^+^ uNK cells. Supporting this finding, lower expression of DNAM1, an activation marker associated with NK cell education^36^, was also found on uNK cells in PCOS-like mice (Figure 3d and Figure S3c). Like NKG2A, this decreased expression of DNAM1 was proportionally constrained to the CD27^+^CD11b^+^ uNK cells (Figure 3e) but was reduced numerically due to fewer DNAM1^+^ mature uNK cells (Figure S3d). Finally, the reduced expression of the MHC class I-specific inhibitory receptor NKG2A on uNK cells in PCOS-like mice raised the question of whether non-MHC class I-specific inhibitory receptors also were affected by androgen exposure. Indeed, a small but distinct population of trNK cells expressing PD1 was found in the uterus, which was reduced in androgen-exposed mice (Figure 3f). Notably, none of the other subsets expressed PD1, nor was it seen on NK cells in blood (Figure S3d), consistent with other reports showing that NK cells do not express PD1 at steady state^37,38^. Taken together, these results show an impaired education of uNK cells in PCOS-like mice, which is tied to the stalled maturation.

**Figure 3.**
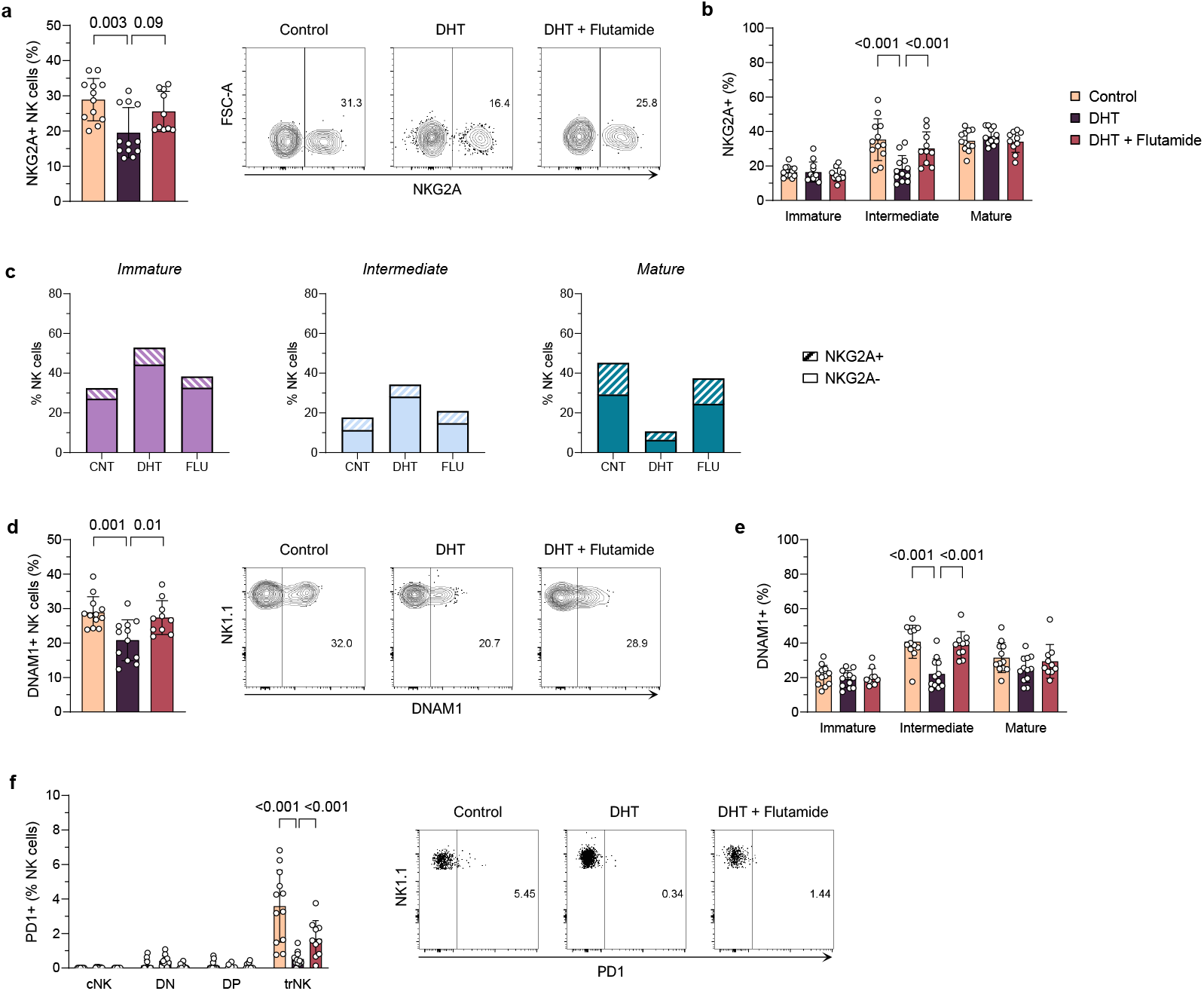
Impaired education of uNK cells in PCOS-like mice. (**a**) Quantification and representative flow cytometry plot of NKG2A+ uNK cells (as % of Lin-NK1.1+ cells). (**b**) Expression of NKG2A on immature (CD27+CD11b-), intermediate (CD27+CD11b+) and mature (CD27-CD11b+) uNK cells. (**c**) Proportions of NKG2A+ uNK cells out of immature, intermediate and mature NK cells, respectively. (**d**) Quantification and representative flow cytometry plot of DNAM1+ uNK cells (as % of Lin-NK1.1+ cells). (**e**) Expression of DNAM1 on immature, intermediate and mature uNK cells. (**f**) Quantification and representative flow cytometry plot of PD1 expression on different subsets of uNK cells. Data are presented as means ± SD.

### Systemic impairment of maturation of NK cells in PCOS-like mice

The pronounced effect of androgen exposure on the maturation of uNK cells led us to ask if this was specific to the uterus or a systemic effect. Therefore, the maturation of NK cells was assessed in other compartments. In keeping with what previously been described^18^, the majority of NK cells in the spleen were mature cNK cells (Figure 4a and Figure S4a). Albeit less evident than in the uterus, a decrease in mature NK cells was observed in the spleen of PCOS-like mice compared to controls and mice co-treated with flutamide. Androgen exposure instead caused a slight increase in intermediate CD27^+^CD11b^+^ NK cells, while no change was observed in immature NK cells. Consistent with an impaired maturation, a lower frequency of the maturation marker KLRG1^+^ NK cells was also found in the spleen of PCOS-like mice (Figure 4b). As in the uterus, androgen exposure caused a small but significant reduction of the education marker NKG2A^+^ NK cells in the spleen (Figure 4c). However, in contrast to the uterus, the reduced NKG2A^+^ expression appeared to be uncoupled from maturation (Figure S4b) and was not associated with any changes in the expression of DNAM1 in the spleen (Figure 4d). The effect of androgen exposure on maturation was mirrored by NK cells in blood (Figure 4e-f). However, no effect on education was observed in circulating NK cells in PCOS-like mice as neither the expression of NKG2A nor DNAM1 was altered in blood (Figure 4g-h). This shows that the impaired maturation of NK cells is a systemic effect in PCOS-like mice, while education is affected in a tissue-specific manner.

**Figure 4.**
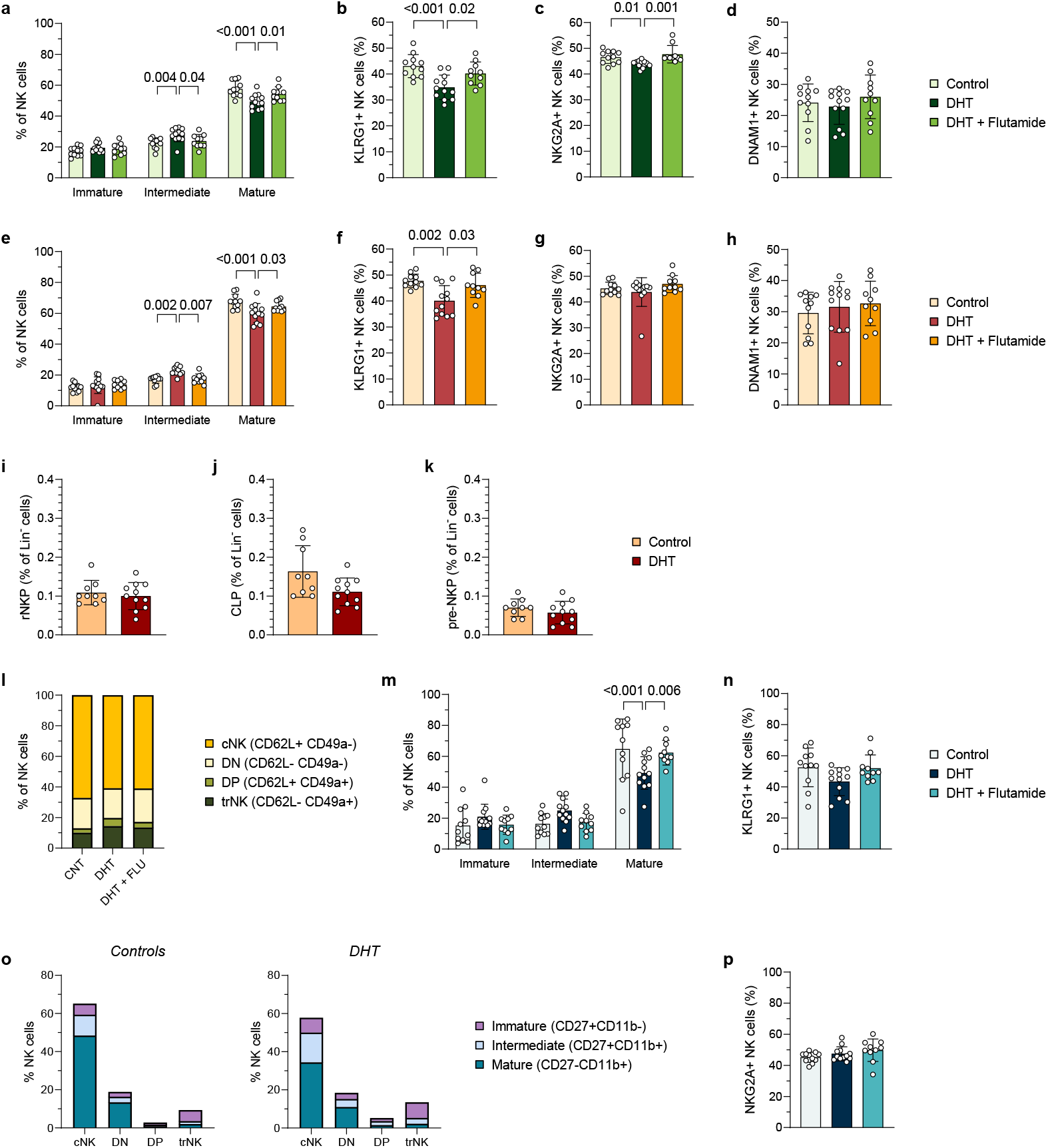
Systemic impairment of maturation of NK cells in PCOS-like mice. (**a**) Frequency of immature (CD27+CD11b-), intermediate (CD27+CD11b+) and mature (CD27-CD11b+) NK cells in spleen. (**b**) Frequency of KLRG1+ NK cells in spleen. (**c**) Frequency of NKG2A+ NK cells in spleen. (**d**) Frequency of DNAM1+ NK cells in spleen. (**e**) Frequency of immature, intermediate and mature NK cells in blood. (**f**) Frequency of KLRG1+ NK cells in blood. (**g**) Frequency of NKG2A+ NK cells in blood. (**h**) Frequency of DNAM1+ NK cells in blood. (**i**) Frequency of real NK cell progenitors (rNKP) in bone marrow. (**c**) Frequency of common lymphoid progenitors (CLP) in bone marrow. (**d**) Frequency of pre-NK cell progenitors (pre-NKP) in bone marrow. (**l**) Proportions of conventional (cNK), double-negative (DN), double-positive (DP) and tissue-resident NK (trNK) cells out of all NK cells in visceral adipose tissue (VAT). (**m**) Frequency of immature, intermediate and mature NK cells in VAT. (**n**) Frequency of KLRG1+ NK cells in VAT. (**o**) Proportions of maturation among subsets in controls and DHT-exposed PCOS-like mice, out of NK cells in VAT. (**p**) Frequency of NKG2A+ NK cells in VAT. Data are presented as means ± SD.

Considering the systemic impact on NK cell maturation, we asked if NK cell progenitors were affected by androgen exposure. Analysis of bone marrow by flow cytometry did not reveal any changes in NK cell progenitors (rNKP, Figure 4i), defined as Lin^−^CD27^+^CD244^+^CD122^+^Flk2^−^. Nor was any change seen in the frequency of common lymphoid progenitors (CLP; Lin^−^CD27^+^CD244^+^CD122^−^Flk2^+^IL-7Rα^+^) or pre-NKPs (Lin^−^CD27^+^CD244^+^CD122^−^Flk2^−^IL-7Rα^+^) in bone marrow from PCOS-like mice (Figure 4j-k).

Lastly, we asked if NK cells in tissues other than secondary lymphoid organs were affected, and if these more closely resembled NK cells in the uterus. Obesity and type-2 diabetes are central comorbidities in the pathology of PCOS^25,27^ and are known to affect fertility and pregnancy outcomes^28–31^. Therefore, the phenotype of NK cells in visceral adipose tissue (VAT) was assessed. In contrast to the uterus, the majority of NK cells in VAT were cNK cells and the proportions of different subsets in VAT from PCOS-like mice were similar to controls and mice co-treated with flutamide (Figure 4l and Figure S4d). Similar to the spleen, the majority of NK cells in VAT were mature, and a slight decrease in mature NK cells was seen in VAT from PCOS-like mice while intermediate and immature NK cells were unchanged (Figure 4m). However, no change in expression of KLRG1 was observed in VAT (Figure 4n), nor did the altered maturation appear to affect any particular subset (Figure 4o). The education of NK cells also remained unaffected by androgen exposure in VAT (Figure 4p and Figure S4e).

To conclude, no uniform systemic effect on NK cells is found in PCOS-like mice. While the maturation of NK cells appears to be inhibited in all tissues, the extent, proportions, and co-expression of KLRG1 differed in all analyzed compartments. Furthermore, androgens do not seem to affect NK cell development upstream of the immature CD27^+^CD11b^−^ subset as progenitors in the bone marrow were unchanged. Only in the uterus was education clearly tied to NK cell maturation; this was not found in any other compartment. Notably, no effect was seen on education of circulating NK cells or in VAT. This demonstrates that NK cells in the uterus should be considered a discrete entity that is distinctively altered by hyperandrogenism.

### Implantation failure coincides with phenotypic alterations of uNK cells in PCOS-like mice

uNK cells are essential to sustain a healthy pregnancy and are believed to promote implantation^39^. It is therefore plausible that alterations of uNK cells could contribute to reproductive dysfunction in our model. We have previously shown that peripubertal DHT-exposed mice have a decreased pregnancy rate compared to controls and mice co-treated with flutamide^40^. However, whether this is due to implantation failure or early miscarriage has not been determined. To address this, PCOS-like mice were superovulated and mated with a fertile male (Figure 5a). Superovulation was necessary because the estrus cycle of PCOS-like mice is arrested in diestrus^41^, an anovulatory stage. The presence of a copulatory plug designated embryonic day 0.5 (E0.5). Implantation sites were visualized by intravenous injection of Evans Blue at E5.5. This time point was selected to distinguish implantation failure from early miscarriage, as blastocysts implant in the uterine wall on the night of E4 in mice^42^. A noticeable difference in the number of implantation sites was observed between PCOS-like mice and controls (Figure 5b-c). In fact, while a third of control mice displayed implantation sites, none of the PCOS-like mice in the cohort were pregnant at E5.5 (Figure 5d), clearly demonstrating implantation failure in PCOS-like mice.

**Figure 5.**
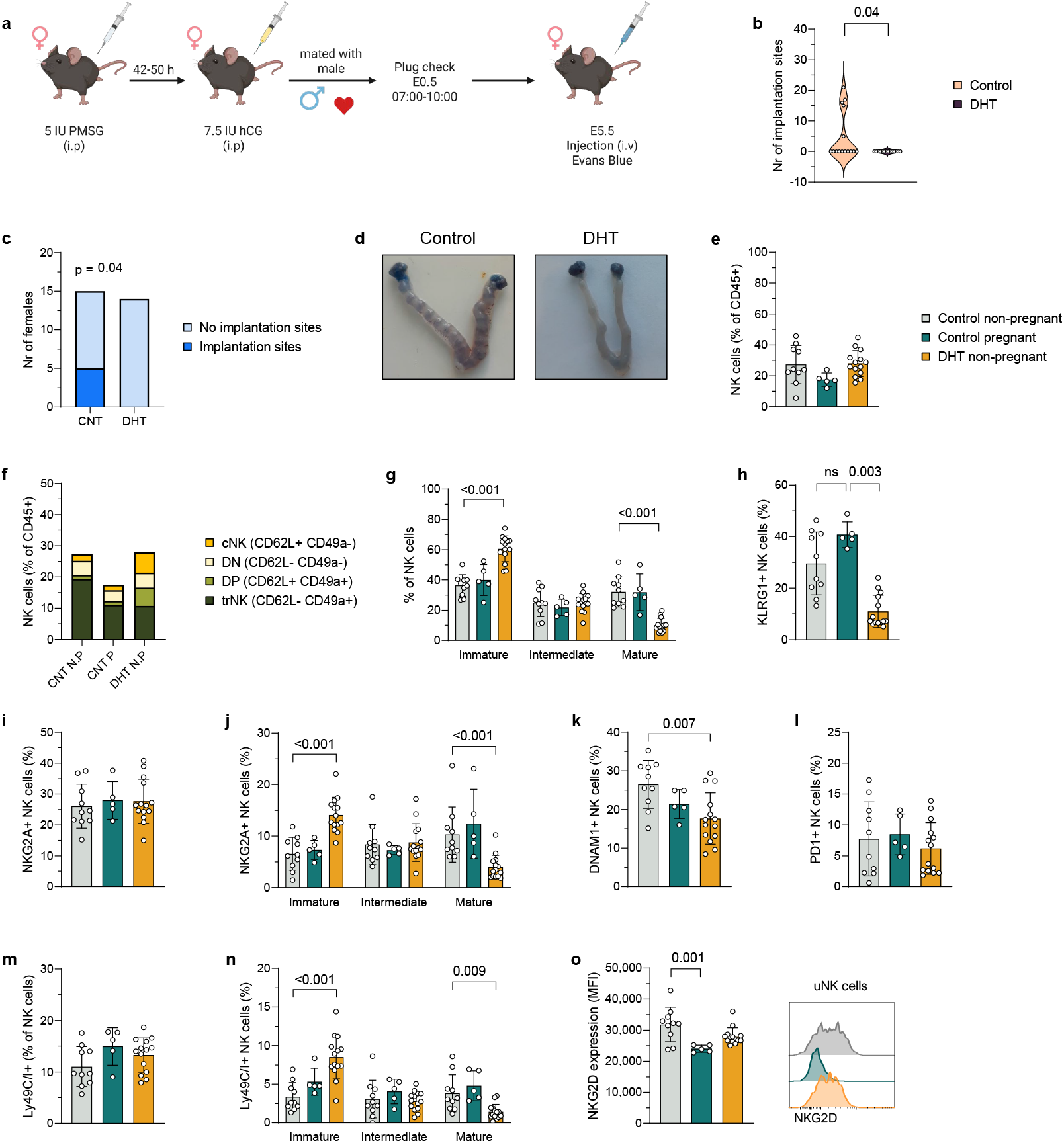
Implantation failure coincide with phenotypic alterations of uNK in PCOS-like mice. (**a**) Experimental design. (**b**) Number of implantation sites at E5.5. (**c**) Frequency of implantation sites at E5.5. (**d**) Representative images of implantation sites at E5.5, visualized by i.v. injection of Evans blue. (**e**) Frequency of uNK cells, out of all CD45+ immune cells in uterus. (**f**) Proportions of conventional (cNK), double-negative (DN), double-positive (DP) and tissue-resident NK (trNK) cells, out of all CD45+ immune cells in uterus. (**g**) Frequency of immature (CD27+CD11b-), intermediate (CD27+CD11b+) and mature (CD27-CD11b+) uNK cells. (**h**) Frequency of KLRG1+ uNK cells. (**i**) Frequency of NKG2A+ uNK cells. (**j**) Expression of NKG2A on immature, intermediate and mature uNK cells. (**k**) Frequency of DNAM1+ uNK cells. (**l**) Frequency of PD1+ uNK cells. (**m**) Frequency of Ly6C/I+ uNK cells. (**n**) Expression of Ly6C/I on immature, intermediate and mature uNK cells. (**o**) Quantification and representative flow cytometry histograms of NKG2D expression on uNK cells. Data are presented as means ± SD.

To investigate which phenotypic changes might contribute to implantation failure, uNK cells were analyzed by the same parameters as previously, now comparing pregnant to non-pregnant controls in addition to PCOS-like mice. Surprisingly, superovulation appeared to have abrogated the increased number of uNK cells in PCOS-like mice (Figure 5e). Even so, the subsets differed proportionally, with a persistently higher frequency of cNK cells and DP uNK cells in PCOS-like mice, while no difference was seen between pregnant and non-pregnant controls (Figure 5f and Figure S5b). The maturation of uNK cells still appeared inhibited, as mature uNK cells constituted only ∼10% of uNK cells in PCOS-like mice, similar to those that had not undergone superovulation (Figure 5g-h and 2g-h). However, no difference was seen on intermediate NK cells following superovulation, while the proportion of immature NK cells had increased further. Notably, the effect on uNK cell education also appeared to be blunted by superovulation as the decrease in NKG2A^+^ uNK cells was abolished (Figure 5i and 3a), as well as the reduced expression of NKG2A on the intermediate CD27^+^CD11b^+^ population (Figure S5b and Figure 3b). However, the number of mature NKG2A^+^ uNK cells was still decreased in PCOS-like mice but compensated by an increase in immature NKG2A^+^ uNK cells which was not seen previously (Figure 5j and Figure S3b). Still, DNAM1^+^ uNK cells remained decreased in PCOS-like mice (Figure 5k). Furthermore, the difference in PD1^+^ uNK cells was also diminished (Figure 5l).

To further analyze the education of uNK cells, the expression of the inhibitory receptors Ly49C and Ly49I, functional homologs of human killer-cell immunoglobulin-like receptors (KIRs), was analyzed. Intriguingly, the expression of Ly49C/I mirrored that of NKG2A, with no overall change in Ly49C/I^+^ uNK cells (Figure 5m) but a decrease in mature Ly49C/I^+^ uNK cells that was compensated by an increase in immature Ly49C/I^+^ uNK cells (Figure 5n). Finally, to better understand the activation state of uNK cells, the activating receptor NKG2D was included in the analysis. Remarkably, a decrease in NKG2D expression was seen on uNK cells in pregnant controls compared to their non-pregnant counterparts and PCOS-like mice (Figure 5o), possibly indicating that lower activation of uNK cells is required during pregnancy.

In conclusion, the distinct phenotypic alterations of uNK cells coincide with implantation failure in PCOS-like mice. Superovulation blunted some of the effects of androgen exposure, possibly due to changes in the hormonal environment. Even so, the major effects on maturation and education of uNK cells persisted. Among the analyzed parameters, only NKG2D was changed in pregnant dams, which could indicate that alterations of uNK cell activation could be of importance in early pregnancy.

## Discussion

Despite the critical role of uNK cells in reproductive function, it is largely unknown how they are affected in PCOS, a syndrome strongly associated with reproductive comorbidities^1,2^. Here, we demonstrate that androgen exposure severely impacts the phenotype of uNK cells and is associated with implantation failure in a well-established peripubertal PCOS-like mouse model. Androgen exposure, mimicking hyperandrogenism, the key diagnostic criteria in PCOS, altered the composition, maturation and education of uNK cells, in an evidently linked manner. First, we show that the previously described expansion of uNK cells in PCOS-like mice^33^ stems from an influx of cNK cells and an increase in CD62L^−^ CD49a^−^ double-negative uNK cells. Recent parabiosis experiments have shown that NK cells from circulation convert into trNK cells in the uterus of normal cycling mice^23^. They also demonstrated that the infiltration and subsequent expansion of trNK cells in the uterus was regulated by progesterone. However, the mechanisms regulating the migration of cNK cells to the uterus remain unknown. How androgen exposure triggers the excessive recruitment of cNK cells to the uterus of PCOS-like mice thus warrants further investigation.

Continuous differentiation and replenishment of uNK cells by circulating NK cells has been demonstrated in humans, where uNK cell differentiation is characterized by the acquisition of KIRs and CD39 in response to IL-15^20^. IL-15 is an important cytokine for the differentiation and survival of NK cells and is produced by endometrial stromal cells when stimulated with progesterone^43–45^. IL-15 also promotes the proliferation of uNK cells, as indicated by higher expression of Ki67 during the secretory phase, but not of cNK cells in the endometrium^20^. This local proliferation was shown to be restricted to undifferentiated KIR^−^ CD39^−^ uNK cells. It is tempting to speculate that the CD62L^+^ CD49a^+^ double-positive (DP) population could constitute a corresponding intermediate population in the uterus of mice, where cNK cells upregulate CD49a before downregulating CD62L during differentiation into trNK cells. We could speculate that the exaggerated influx of cNK cells is coupled with an inability to convert to trNK cells, or even a compensatory mechanism for this reduced ability. The lower expression of TGF-β1 in the uterus of PCOS-like mice would support this hypothesis, as autocrine signaling with TGF-β1 has been shown to be essential for the transformation of cNK cells into trNK cells^22^. An inability to convert to trNK cells could be linked to the disrupted maturation of NK cells seen in PCOS-like mice, as the expression pattern of CD49a to CD27 suggests a relationship between these markers and was found to differ between PCOS-like mice and controls. An association between maturation and tissue residency would be in line with previous work showing that immature cNK cells are more disposed to convert into trNK cells when stimulated with TGF-β1 than mature cNK cells^22^. On this basis, the possibility of DP uNK cells as an intermediate state of cNK cells differentiating into trNK cells would be likely, as this population exclusively consisted of immature NK cells in the uterus of control mice.

The CD62L^−^ CD49a^−^ double-negative (DN) population of uNK cells was almost exclusively mature in the uterus of control mice, making this subset unlikely to represent an intermediate state of transitioning cNK cells. To our knowledge, the DN uNK cell population has not previously been described. Phenotypically, the DN subset resembled cNK cells more than trNK cells, given their maturation and lack of CD69 expression in controls. Interestingly, not only was this subset vastly increased in PCOS-like mice, but it had also acquired expression of CD69. A small but distinct proportion of cNK cells in the uterus of PCOS-like mice also upregulated CD69, which is noteworthy, as no difference in CD69 expression was observed in blood. This suggests that the upregulation of CD69 is coupled to the infiltration of cNK cells into the uterine tissue. Moreover, this effect coincided with a decreased frequency of CD69^+^ trNK cells, which leads to the question of whether the acquisition of CD69 by DN and cNK cells is a compensatory mechanism. While human uNK cells express high levels of CD69^46,47^, the function of CD69 in reproduction has not yet been described. However, galectin-1, one of the ligands for CD69, has been shown to be pivotal for a successful pregnancy, as knockout of *Lgals1*, the gene encoding galectin-1, renders mice more susceptible to fetal loss^48^. External administration of galectin-1 rescued a stress-induced model of pregnancy loss and upregulated the expression of TGF-β1, plausibly linking CD69 to the maintenance of trNK cells or the reprogramming of cNK cells into tissue residency^22^. Furthermore, galectin-1 increases in the human endometrium during the secretory phase and remains high in the decidua^49^. The expression has been shown to be regulated by progesterone^50^, possibly linking galectin-1 to the expansion of trNK cells in the uterus^20,23,51^. While galectin-1 expression peaks at gestational day 5, when implantation occurs in mice^50^, no difference in the number of implantations sites was reported in *Lgals1*^−/−^ mice compared to controls^48^. It is therefore uncertain whether altered expression of CD69 on uNK cells could contribute to the implantation failure in PCOS-like mice. Even so, single nuclei RNA sequencing data of endometrium from women with PCOS show a downregulation of *LGALS1* on cycling stromal cells^32^, which could imply clinical relevance of altered CD69 expression on uNK cells in our model.

Education of NK cells is a process that fine-tunes their responsiveness, coordinated by activating and inhibitory receptors^52^. This process is crucial for fetomaternal tolerance and regulates important functions of uNK cells, such as their ability to promote the formation of spiral arteries^15,35^. We demonstrate that uNK cells in PCOS-like mice have an impaired education, evident by a reduced frequency of uNK cells expressing the inhibitory receptor NKG2A. In contrast to circulating NK cells, human uNK cells express high levels of NKG2A^53,54^. The CD94–NKG2A complex binds HLA-E^55^, which is expressed by trophoblasts, and thereby inhibiting NK cell cytotoxicity^56^. This interaction has also been shown to promote appropriate vascularization. The non-classical HLA-E requires peptides provided by classical HLA-A, -B or -C to be functional^57,58^. Women with an HLA-B dimorphism, leading to a non-functional interaction between HLA-E and NKG2A, have a higher risk of pre-eclampsia^15^. Furthermore, NKG2A-deficient dams display improper vascularization during pregnancy, which is associated with aberrant placental gene expression and fetal growth-restriction^15^. NKG2A-deficiency does not affect litter size and is therefore not likely to be the underlying cause of implantation failure in PCOS-like mice. However, the reduced frequency of NKG2A^+^ uNK cells in PCOS-like mice is an interesting finding, as women with PCOS have an increased risk of pre-eclampsia and warrants further investigation.

The impaired education of uNK cells in PCOS-like mice was evidently linked to maturation, as the reduced frequency of NKG2A^+^ uNK cells clearly was due to a decrease of mature uNK cells expressing NKG2A. Interestingly, superovulation appear to trigger an increase of NKG2A^+^ immature uNK cells in PCOS-like mice, rendering the overall frequency of NKG2A^+^ uNK cells equal to that of controls. This expression pattern was mirrored by Ly49C and Ly49I, inhibitory receptors and functional homologs of human killer-cell immunoglobulin-like receptors (KIRs), underscoring the link between maturation and education as well as the tissue specific impairment of NK cell education in the uterus of PCOS-like mice. This demonstrates once again the influence of sex hormones on uNK cells, as it is likely that changes to the uNK cell population in PCOS-like mice following superovulation is due to hormonal changes in the uterine environment. Collectively, the results highlight the difficulty of treating immune-related reproductive complications, considering the dynamic properties of uNK cells, and more importantly, their dependency on the local tissue environment. Indeed, while there was a systemic effect on the maturation of NK cells in PCOS-like mice, NK cells in circulation, secondary lymphoid tissues and adipose tissue were vastly differently affected by androgen exposure compared to tissue-resident uNK cells.

To conclude, our preclinical model suggests that hyperandrogenism can have a considerable impact on the function of uNK cells. These findings clearly indicate that a disruption of uNK cells contributes to the endometrial dysfunction observed in PCOS and may underly the reproductive comorbidities associated with the syndrome, although several aspects remain to be explored, including the function of DN uNK cells, identification of the human counterpart, the role of CD69 in reproductive function, and the exact mechanism underlying implantation failure in our model.

## Methods

### Animals and implants for inducing a PCOS-like mouse model

C57BL/6JRj females were purchased from Janvier Labs and housed in a temperature-controlled environment under a 12-hour light/dark cycle. Mice had ad libitum access to water and a standard chow diet (CRM (P), SAFE Diets). The peripubertal DHT-exposed PCOS-like mouse model was induced as previously described^59^. Briefly, a continuously releasing DHT implant was inserted subcutaneously in the neck region of 28 to 29-day old females under anesthesia with 1.5-2% isoflurane (Vetflurane, Virbac). Implants were made by inserting an average of 3.2 mg DHT (5α-Androstan-17β-ol-3-one, ≥97.5%, Sigma-Aldrich) into silastic tubes trimmed to a total length of 5 mm. Empty, blank implants were inserted in control mice. Mice co-treated with flutamide were implanted with a 25 mg continuously releasing flutamide pellet (NA-152, Innovative Research of America) in addition to the DHT implant. All animal experiments were approved by the Stockholm Ethical Committee for animal research (18003-2023) in accordance with the Swedish Board of Agriculture’s regulations and recommendations (SJVFS 2019:9) and the European Parliament’s directive on the protection of animals used for scientific purposes (2010/63/EU). Animal care and procedures were performed in accordance with the guidelines of the Federation of European Laboratory Animal Science Associations (FELASA) and overseen by Comparative Medicine Biomedicum at the Karolinska Institutet in Stockholm, Sweden.

### Tissue collection

Mice were sacrificed during diestrus, determined by vaginal cytology as previously described^60^, at 16-19 weeks of age. Blood was drawn by cardiac puncture under anesthesia with 3% isoflurane (Vetflurane, Virbac) and collected into EDTA-coated tubes (Sarstedt). For flow cytometric analysis, uteri and perigonadal VAT were collected in RPMI-1640 (Sigma-Aldrich) supplemented with 2% fetal bovine serum (FBS, Gibco™). Spleen were collected in Ca^2+^ and Mg^2+^ free DPBS (Sigma-Aldrich). All samples were kept on ice. For gene expression analysis, a small piece from one uterine horn was snap-frozen in liquid nitrogen and stored at −80°C.

### Flow cytometry

Uteri were cut and dissociated by gentle shaking for 20-25 min at 37°C, in a digestion mix containing collagenase type I (1 mg/ml, Sigma-Aldrich), 0.8U DNase I (Sigma-Aldrich) and 2% FBS (Gibco™) in RPMI-1640 (Sigma-Aldrich). VAT was minced using fine scissors and gently shaken at 37°C for 20-30 min in a digestion mix of collagenase type IV (1 mg/ml, Sigma-Aldrich), 2% FBS (Gibco™) in RPMI-1640 (Sigma-Aldrich). Bone marrow from one femur was collected into flow cytometry buffer (2% FBS and 2 mM EDTA in PBS) by centrifugation at 500G for 5 min at +4°C. Bones were flushed with flow cytometry buffer to retrieve remaining cells. Single-cell suspensions were obtained by passing samples through a 100μm strainer. Spleen were directly passed through a 100μm strainer. Cells were washed in flow cytometry buffer. Hemolysis of erythrocytes was done in VAT, blood and spleen (0.16 M NH4Cl, 0.13 M EDTA and 12 mM NaHCO3 in H2O) followed by additional wash. Cells were incubated in Fc-block (anti-mouse CD16/32, clone 2.4G2, BD Biosciences) to prevent unspecific binding. Dead cells were stained using Zombie NIR™ Fixable Viability Kit (BioLegend). The following fluorochrome-conjugated antibodies were used to detect cell surface markers in uterus, VAT, spleen and blood: CD45 (30-F11, Thermo Fisher Scientific), CD3e (17A2, BioLegend or 500A2, BD Biosciences), CD8a (53-6.7, BioLegend), CD19 (1D3, BioLegend), NK-1.1 (PK136, BD Biosciences), CD49a (Ha31/8, BD Biosciences), CD62L (MEL-14 BioLegend), CXCR6 (SA051D1, BioLegend), CD49b (DX5, Invitrogen), CD69 (H1.2F3, eBioscience), CD11b (M1/70, BD Biosciences), CD27 (LG.7F9, Thermo Fisher Scientific), KLRG1 (2F1, BioLegend), NKG2A (20d5, BD Biosciences), DNAM1 (TX42.1, BD Biosciences), PD1 (J43, BD Biosciences), Ly49C/I (5E6, BD Biosciences), NKG2D (CX5, BioLegend). The following fluorochrome-conjugated antibodies were used to detect cell surface markers in bone marrow: 2B4 (eBio244F4, Thermo Fisher Scientific), CD11b (M1/70, BD Biosciences), CD122 (TM-β1, BD Biosciences), CD127 (A7R34, BioLegend), CD19 (1D3, BioLegend), CD27 (LG.7F9, Thermo Fisher Scientific), CD3 (17A2, BioLegend), c-Kit (2B8, BD Biosciences), FLT3 (A2F10.1, BD Biosciences), Ly-6D (49-H4, BioLegend), NK-1.1 (PK136, BD Biosciences), NKp46 (29A1.4, BD Biosciences). Cells were analyzed on an ID7000™ Spectral Cell Analyzer (Sony Biotechnology) with 355, 405, 488, 561 and 640nm lasers. Data was processed on ID7000 Software v2.2 (Sony) and further analyzed with FlowJo™ v10.10 Software (BD Life Sciences).

### Gene expression analysis

Total RNA was isolated from uteri using TRI reagent (T9424, Sigma). A High-Capacity RNA-to-cDNA kit (4387406, Applied Biosystems, Carlsbad, California, USA) was used for reversed transcription. Quantitative real-time PCR was performed in a quantflex 6 Real-Time PCR system thermal cycler with SYBR Green PCR Master Mix (both Applied Biosystems). Primer sequences for gene expression analysis of *TGF-β1* were F: ATTCCTGGCGTTACCTTGG and R: AGCCCTGTATTCCGTCTCCT. *Gapdh* (F: AGGTCGGTGTGAACGGATTTG, R: GGGGTCGTTGATGGCAA) was selected as the endogenous control. Data was analyzed using the ΔCt method.

### Single nuclei RNA seq

Single-nuclei RNA sequencing data from endometrial immune cells were obtained from a previously published study (PECA study^32^) and updated to Seurat v5^61^. The eNK cell populations (eNK 1-3) were a subset from the endometrial immune cell atlas, keeping only samples derived from women with and without PCOS at baseline (Figure S2k). Quality control metrics including UMI counts, detected genes, and mitochondrial/ribosomal/hemoglobin percentages were evaluated for each sample. Samples with fewer than 100 nuclei were excluded, resulting in 14 samples (5 controls, 9 PCOS) comprising 3,624 eNK nuclei for analysis. Integration of the samples was performed using Seurat v5’s IntegrateLayers with canonical correlation analysis (CCA) after normalization with LogNormalize and variable feature selection (n = 2,000) using variance-stabilizing transformation (VST)^61^. The eNK cell populations were validated using previously established markers^51^: *NCAM1* (CD56), *FGFBP2* (CD16-negative), *ENTPD1* (CD39), *ITGB2* (CD18), *ITGAE* (CD103), and *CD160* (Figure S2k). TGF-β signaling was examined by gene expression of *TGFB1, TGFBR1*, and *TGFBR2*. Gene co-expression was analyzed by FeaturePlot with blend mode (blend threshold = 0.25) to identify nuclei co-expressing TGF-β ligand-receptor pairs.

### Superovulation and visualization of implantation sites

Mice from all groups were superovulated 4-8 weeks after model induction with 5 IU pregnant mare’s serum gonadotropin (PMSG; Folligon, MSD Animal Health Care) followed by 7.5 IU human chorionic gonadotropin (hCG; Chorulon Vet. 1500 IU, MSD Animal Health Care) 48 hours later. Females were then mated overnight with a normal, fertile male immediately after hCG injection (2 females were paired with 1 male). The presence of a copulatory plug the following morning designated embryonic day 0.5 (E0.5). At E5.5, mice were anesthetized with 3% isoflurane (Vetflurane, Virbac) and 0.1 mL of a 1% solution of Evans Blue was injected intravenously into the tail. Implantation sites were clearly visible as blue bands upon sacrifice 3 minutes after injection (see Figure 5d).

### Statistical analysis

Sample size was predetermined solely based on previously reported assessments^33,40,41,62^. No statistical power calculations were performed. Mice were arbitrarily assigned to experimental groups without formal randomization, and investigators were not formally blinded during the experiment. All data are presented as mean ± SD, with one animal considered an experimental unit. Normality was assessed by Shapiro-Wilk test. Independent, normally distributed data with equal variance among groups were analyzed by one-way ANOVA and Dunnett’s multiple comparison test. For data with normal distribution but unequal variance, Brown-Forsythe and Welch ANOVA tests with Dunnett’s multiple comparison test was used. Kruskal-Wallis and Dunn’s multiple comparison test were used to analyze data that did not follow a normal distribution. Two-way ANOVA and Dunnett’s multiple comparison test were used for interacting variables. Relative RNA expression by qPCR was analyzed by two-tailed Welch’s test. Implantation sites and proportion of pregnant dams were calculated by two-tailed Mann-Whitney test and Fisher’s exact test respectively. P ≤ 0.05 was considered statistically significant. Statistical analyses were performed using GraphPad Prism 10 (GraphPad Software).

## Data availability

The source data supporting the findings of this study are provided in BioStudies and will be publicly available upon publication of this paper under accession number: S-BSST2694.

## Code availability

The code used to generate and analyze the snRNA-seq is available at: github.com/ReproductiveEndocrinologyMetabolism/PCOS_eNK_Implantation_Analysis

## Graphics

Figures 1a, 5a were created with BioRender.com.

## Author contribution

Conceptualization: ST; ESV. Methodology: ST; ESV; MHJ; BJC. Investigation: ST; HL; AZ; CG; EL; GE. Data acquisition, analysis, and visualization: ST; MHJ; BJC; CG; ESV. Project administration: ST; ESV. Supervision: ESV; AB; MHJ; BJC. Wrote the manuscript: ST; ESV. All authors: reviewing and editing manuscript. Funding acquisition: ESV.

## Acknowledgement

We thank the Biomedicum Flow cytometry Core facility (Karolinska Institutet), supported by KI/SLL, for providing access to equipment, technical expertise and scientific input. This work is supported by Swedish Medical Research Council: project no. 2022-00550 (ESV); Distinguished Investigator Grant – Endocrinology and Metabolism, Novo Nordisk Foundation: NNF22OC0072904 and project grant: NNF25OC0104784 (ESV); Diabetes Foundation: DIA2022-708 and DIA2024-860 (ESV); Karolinska Institutet KID funding: 2020-00990 (ESV).

