## Supplemental figures for "Implantation failure linked to altered uterine NK cells in PCOS-like mice"

### Supplemental information

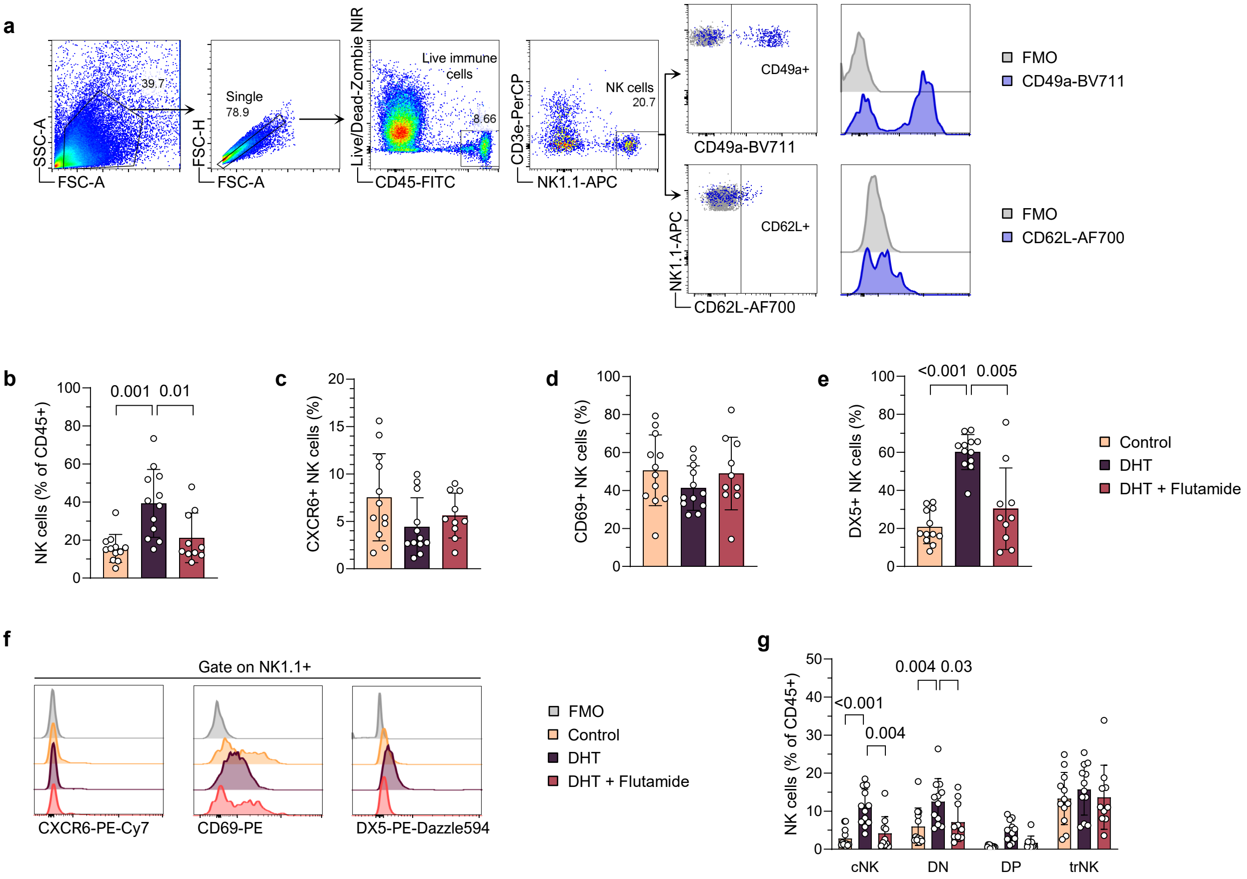


**Figure S1. Augmented population of uNK cells in PCOS-like mice is due to an increased infiltration of cNK cells.** (**a**) Gating strategy. (**b**) Frequency uNK cells, out of all CD45+ immune cells in uterus. (**c**) Frequency CXCR6+ uNK cells (as % of Lin-NK1.1+ cells). (**d**) Frequency CD69+ uNK cells (as % of Lin-NK1.1+ cells). (**e**) Frequency DX5+ uNK cells (as % of Lin-NK1.1+ cells). (**f**) FMOs and representative flow cytometry histograms of CXCR6, CD69 and DX5 expression on uNK cells. (**g**) Frequency of conventional (cNK; CD62L+CD49a-), double-negative (DN; CD62L-CD49a-), double-positive (DP; CD62L+CD49a+) and tissue-resident (trNK; CD62L-CD49a+) uNK cells. Data are presented as means ± SD.

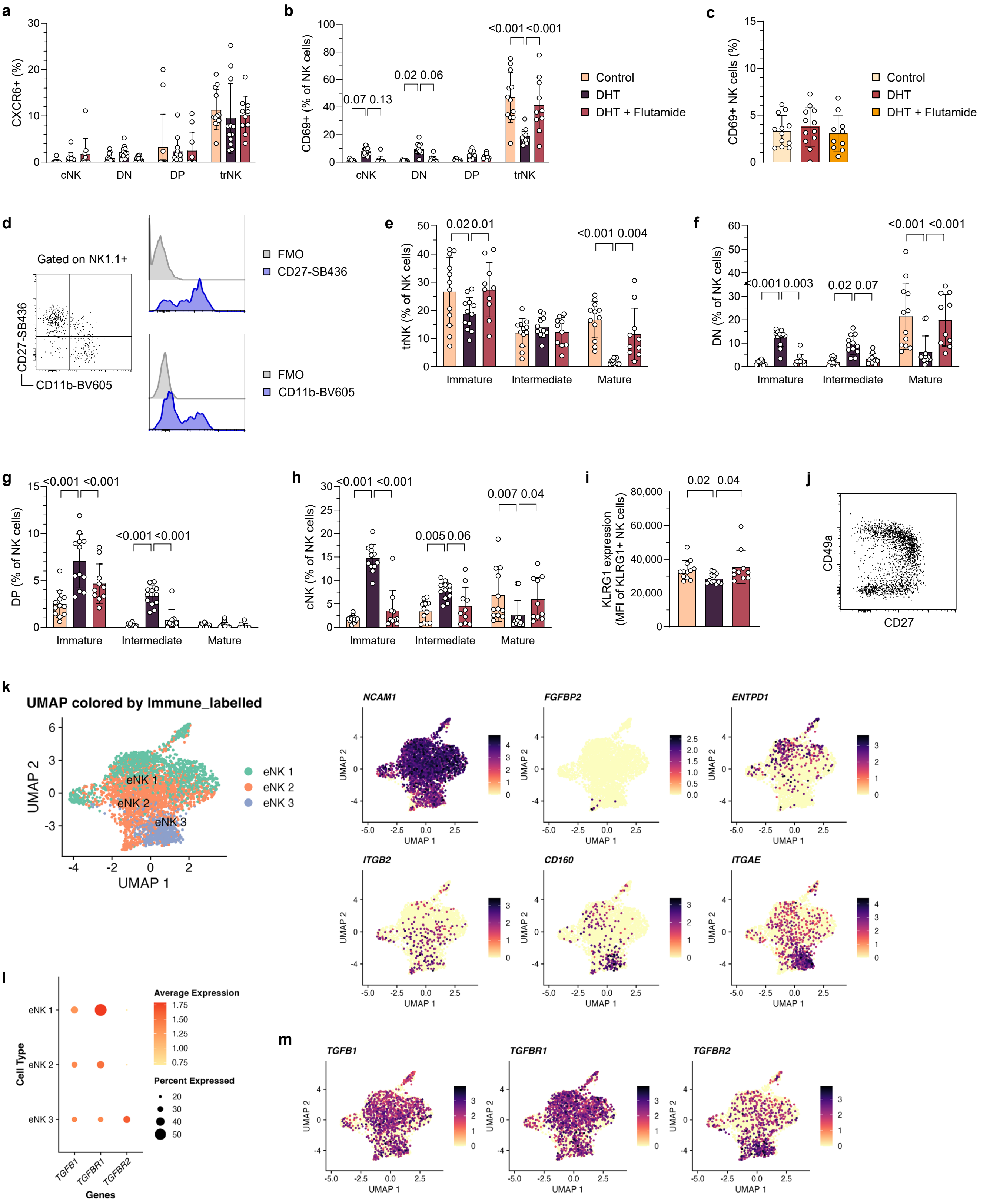


**Figure S2. Distinct phenotypic alterations of uNK subsets in PCOS-like mice**

Figure legend continues at next page

**Figure S2. Distinct phenotypic alterations of uNK subsets in PCOS-like mice** (**a**) CXCR6 expression on conventional (cNK; CD62L+CD49a-), double- negative (DN; CD62L-CD49a-), double-positive (DP; CD62L+CD49a+) and tissue-resident (trNK; CD62L-CD49a+) uNK cells. (**b**) Frequency of CD69+ cNK, DN, DP and trNK cells. (**c**) Frequency of CD69+ NK cells in blood (as % of Lin-NK1.1+ cells). (**d**) Representative flow cytometry plots and FMOs of CD27 and CD11b in uterus. (**e**) Frequency of immature (CD27+CD11b-), intermediate (CD27+CD11b+) and mature (CD27-CD11b+) trNK cells in uterus. (**f**) Frequency of immature, intermediate and mature DN uNK cells. (**g**) Frequency of immature, intermediate and mature DP uNK cells. (**h**) Frequency of immature, intermediate and mature cNK cells in uterus. (**i**) Expression of the maturation marker KLRG1 uNK cells. (**j**) Expression pattern of CD49a and CD27 on uNK cells. (**k**) UMAP projections of integrated snRNA-seq data of endometrial NK (eNK) cells from women with and without PCOS, with annotations of endometrial eNK cell subpopulations (eNK1, eNK2, eNK3; left) and expression gene markers for validation (right): *NCAM1* (CD56), *FGFBP2* (CD16-negative), *ENTPD1* (CD39), *ITGB2* (CD18), *ITGAE* (CD103), and *CD160*. (**l**) Dot plot displaying expression of TGF-β (TGFB1), TGF-β receptor 1 (TGFBR1), and TGF-β receptor 2 (TGFBR2) on eNK cell subpopulations (eNK1-3). (**m**) UMAP projections of integrated snRNA-seq data displaying expression of TGF-β (TGFB1), TGF-β receptor 1 (TGFBR1), and TGF-β receptor 2 (TGFBR2) on eNK cells. Data are presented as means ± SD.

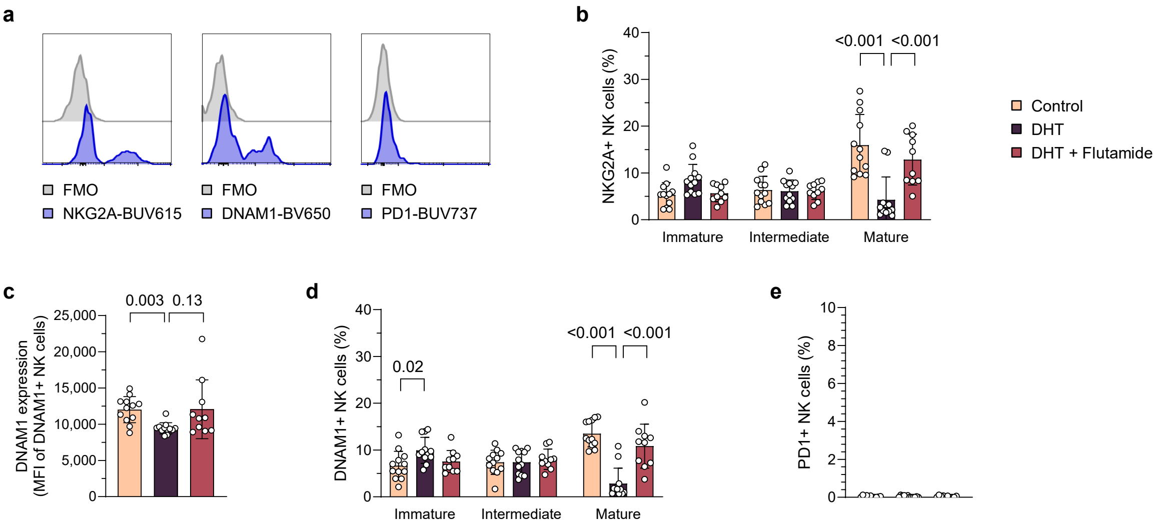


**Figure S3. Impaired education of uNK cells in PCOS-like mice** (**a**) FMOs of NKG2A, DNAM1 and PD1 on uNK cells. (**b**) Frequency of NKG2A+ immature (CD27+CD11b-), intermediate (CD27+CD11b+) and mature (CD27-CD11b+) uNK cells. (**c**) Expression of DNAM1 on uNK cells. (**d**) Frequency of DNAM1+ immature, intermediate and mature uNK cells. (**e**) Frequency of PD1+ NK cells in blood. Data are presented as means ± SD.

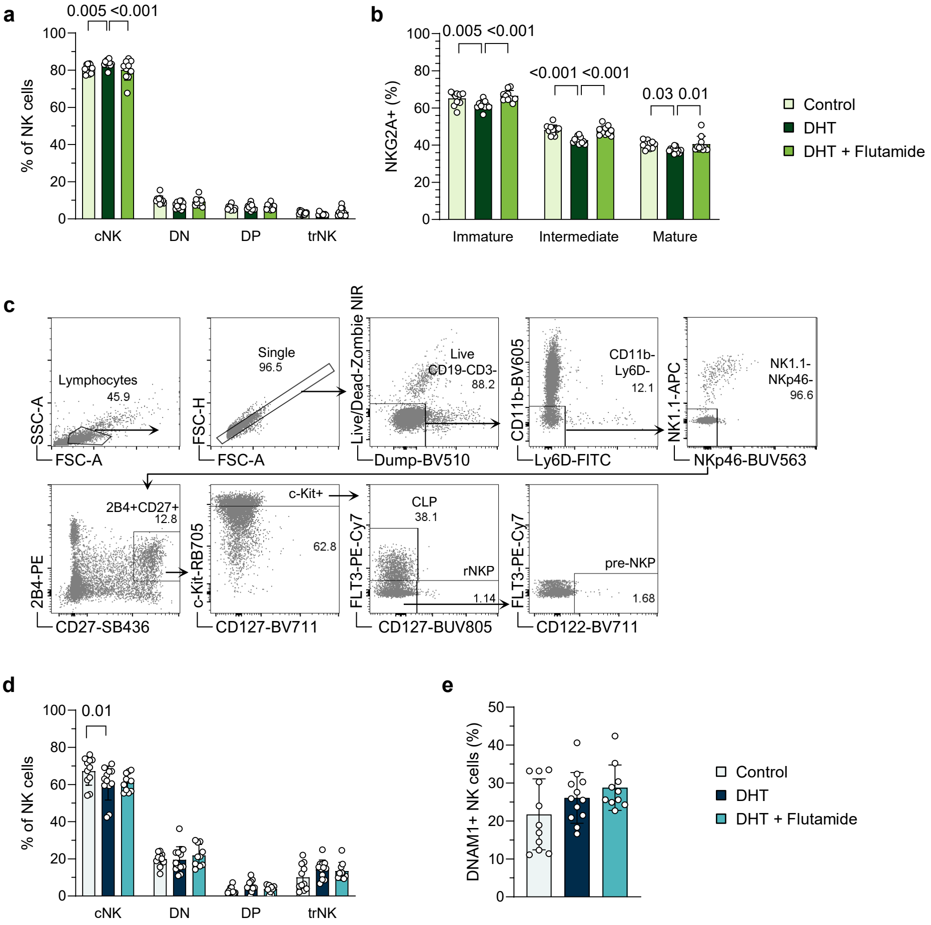


**Figure S4. Systemic impairment of maturation of NK cells in PCOS-like mice** (**a**) Frequency of conventional (cNK; CD62L+CD49a-), double-negative (DN; CD62L-CD49a-), double-positive (DP; CD62L+CD49a+) and tissue-resident NK (trNK; CD62L-CD49a+) cells in spleen. (**b**) Frequency of NKG2A+ immature (CD27+CD11b-), intermediate (CD27+CD11b+) and mature (CD27-CD11b+) NK cells in spleen. (**c**) Gating strategy of real NK cell progenitors (rNKP), common lymphoid progenitors (CLP) and pre-NK cell progenitors (pre-NKP) in bone marrow. (**d**) Frequency of conventional (cNK), double-negative (DN), double-positive (DP) and tissue-resident NK (trNK) cells out of all NK cells in visceral adipose tissue (VAT). (**e**) Frequency of DNAM1+ NK cells in VAT (as % of Lin-NK1.1+ cells). Data are presented as means ± SD.

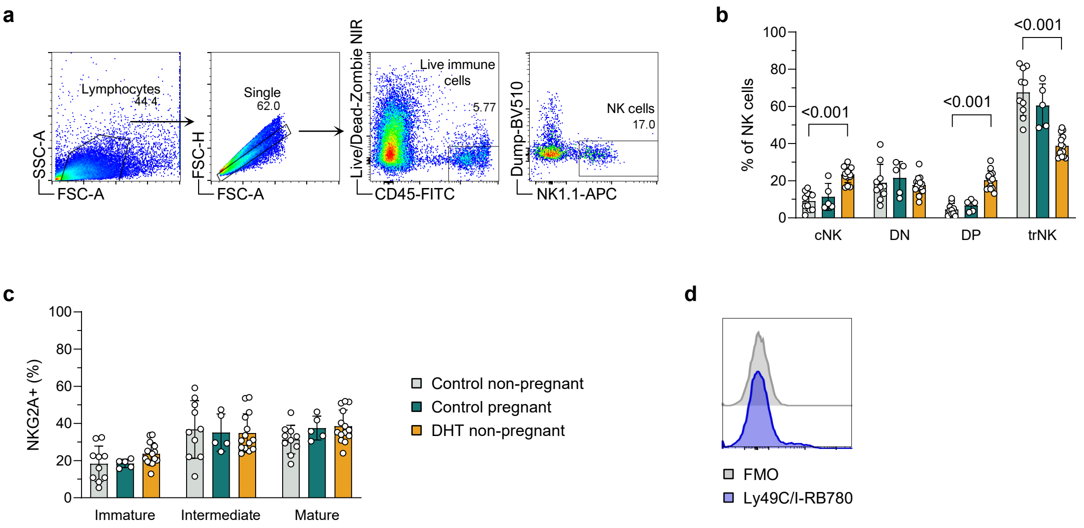


**Figure S5. Implantation failure coincide with phenotypic alterations of uNK in PCOS-like mice** (**a**) Gating strategy. (**b**) Frequency of conventional (cNK; CD62L+CD49a-), double-negative (DN; CD62L-CD49a-), double-positive (DP; CD62L+CD49a+) and tissue-resident NK (trNK; CD62L-CD49a+) cells in uterus. (**c**) Frequency of NKG2A+ immature (CD27+CD11b-), intermediate (CD27+CD11b+) and mature (CD27-CD11b+) uNK cells. (**d**) FMO of Ly49C/I on uNK cells. Data are presented as means ± SD.
